# The Role of Toll-like receptor 4 in respiratory syncytial virus replication, interferon lambda 1 induction, and chemokine responses

**DOI:** 10.1101/404384

**Authors:** Lindsay Broadbent, Jonathon D. Coey, Michael D. Shields, Ultan F. Power

## Abstract

Respiratory syncytial virus (RSV) infection is the leading cause of severe lower respiratory tract infections (LRTI) in infants worldwide. The immune responses to RSV infection are implicated in RSV pathogenesis but RSV immunopathogenesis in humans remains poorly understood. We previously demonstrated that IFN-λ1 is the principle interferon induced following RSV infection of infants and well-differentiated primary pediatric bronchial epithelial cells (WD-PBECs). Interestingly, RSV F interacts with the TLR4/CD14/MD2 complex to initiate secretion of pro-inflammatory cytokines, while TLR4 stimulation with house dust mite induces IFN-λ1 production. However, the role of TLR4 in RSV infection and concomitant IFN-λ1 induction remains unclear. Using our RSV/WD-PBEC infection model, we found that CLI-095 inhibition of TLR4 resulted in significantly reduced viral growth kinetics, and secretion of IFN-λ1 and pro-inflammatory chemokines. To elucidate specific TLR4 signalling intermediates implicated in virus replication and innate immune responses we selected 4 inhibitors, including LY294002, U0126, SB203580 and JSH-23. SB203580, a p38 MAPK inhibitor, reduced both viral growth kinetics and IFN-λ1 secretion, while JSH-23, an NF-κB inhibitor, reduced IFN-λ1 secretion without affecting virus growth kinetics. Our data indicate that TLR4 plays a role in RSV entry and/or replication and IFN-λ1 induction following RSV infection is mediated, in part, by TLR4 signalling through NF- κB and/or p38 MAPK. Therefore, targeting TLR4 or downstream effector proteins could present novel treatment strategies against RSV.

**Importance:** The role of TLR4 in RSV infection and IFN-λ1 induction is controversial. Using our WD-PBEC model, which replicates many hallmarks of RSV infection *in vivo*, we demonstrated that the TLR4 pathway is involved in both RSV infection and/or replication and the concomitant induction of IFN-λ1 and other pro-inflammatory cytokines. Increasing our understanding of the role of TLR4 in RSV immunopathogenesis may lead to the development of novel RSV therapeutics.

## Introduction

Respiratory syncytial virus (RSV) is responsible for approximately 3.4 million hospitalisations of infants each year and is the primary cause of severe lower respiratory tract infections (LRTI) in children worldwide(1,2). There are currently no RSV vaccines or specific therapeutics available. The innate immune system provides an important first line of defence against RSV disease. Innate immune signalling pathways are initiated by the recognition of pathogen associated molecular patterns (PAMPs) by cell surface or intracellular pathogen recognition receptors (PRRs), such as Toll-like receptors (TLRs). Some TLRs are implicated in activating innate immune responses following RSV infection. In particular, RSV F was shown to interact with the TLR4/CD14/MD2 complex, thereby initiating a signalling cascade leading to the induction of pro-inflammatory cytokines(3-5).

In infants hospitalised with RSV bronchiolitis TLR4 was shown to be upregulated on peripheral blood monocytes during the acute phase of the disease(6). However, the role of TLR4 in RSV infection and immunopathogenesis remains controversial. Some studies in mice demonstrated a TLR4-dependent innate immune response following RSV infection(3,4,7). Other studies in cell lines expressing human TLR4, however, reported that TLR4 did not play a significant role in RSV entry or NF-κB activation(8). The p38 MAPK pathway, which is downstream of TLR4, was implicated in RSV replication and was also shown to be involved in the expression of TLR4 near the site of infection. Furthermore, inhibition of TLR4 resulted in a decrease in p38 MAPK activity(9).

We previously described well-differentiated primary pediatric bronchial epithelial cell (WD-PBEC) cultures that replicated many morphological and physiological hallmarks of bronchial epithelium *in vivo*, including ciliated epithelium and mucus producing goblet cells(10). Airway epithelium is the primary target for RSV infection *in vivo* (11). Importantly, we also demonstrated that RSV infection of WD-PBECs reproduces several hallmarks of RSV infection in infants. To date, much of the research to elucidate pathways leading to the induction of pro-inflammatory chemokines and cytokines following RSV infection has focused on the use of continuous cell lines or semi-permissive animal models. WD-PBECs evidently provide a more biologically relevant model in which to study innate immune responses to RSV infection of the human airway epithelium.

Our aim was to exploit our WD-PBEC model to investigate the role of TLR4 in RSV infection, the induction of type III IFNs, specifically IFN-λ1, and the production of pro-inflammatory chemokines by using specific inhibitors of TLR4 or downstream components of this pathway. CLI-095 is a potent and highly specific inhibitor of TLR4 signalling. It binds to Cys747 in the TLR4 intracellular domain but does not inhibit the binding of ligands to TLR4. CLI-095 binding disrupts the interaction of TLR4 with adaptor molecules, such as MyD88(12). We also exploited inhibitors of NF-κB (JSH- 23), p38MAPK (SB203580), PI3K (LY294002), and MEK1/2 (U0126) to dissect signalling pathways implicated in RSV infection/replication and innate immune response induction.

We demonstrated that the intracellular signalling mediated by the TLR4 complex was involved in RSV infection/replication in WD-PBECs and initiated downstream innate immune responses, including type III IFNs and pro-inflammatory chemokines. p38 MAPK signalling was also implicated in RSV replication. Our data demonstrated that inhibition of TLR4 signalling decreases RSV-induced IFN-λ1 secretion. Furthermore, inhibition of p38 MAPK or NF-κB resulted in a reduction in RSV-induced IFN-λ1 secretion. As NF-κB inhibition did not affect RSV replication kinetics but diminished IFN-λ1 secretion, our data are consistent with an NF-κB-dependent mechanism of IFN-λ1 induction following RSV infection.

## Results

To determine whether TLR4 influences RSV infection, HEK293 cells stably transfected with TLR4 or control HEK293/null cells were infected with RSV A2/eGFP (MOI∼0.1). HEK293 cells do not endogenously express TLR4 (Figure 1A). The extent of eGFP fluorescence in the monolayers was used as a surrogate for the level of RSV infection. The kinetics and peak of eGFP spread throughout the monolayers was significantly higher in HEK293/TLR4 than in the HEK293/null cells, indicating that the presence of TLR4 increased the susceptibility of these cells to RSV infection (Figure 1B). We used CLI-095, a highly selective TLR4 signalling blocker, to investigate if it was the presence of TLR4 on the cell surface or the subsequent signalling cascade that was involved in the increase in RSV infection. CLI-095 does not affect ligand binding to TLR4 but inhibits all downstream signalling following TLR4 activation. CLI-095 (10 µg/mL) treatment of HEK293/TLR4 cells significantly reduced RSV growth kinetics compared to untreated HEK293/TLR4 controls, as evidenced by eGFP fluorescence spread over time. At 48 hpi there was a significant difference between the level of eGFP expression in HEK293/null cells compared to HEK293/TLR4 (p=0.006) and HEK/TLR4 cells compared to HEK293/TLR4 vs HEK293/TLR4 + CLI-095 (p=0.0152). Similarly, at 72 hpi there was a significant difference between HEK293/null cells and HEK293/TLR4 (p<0.0001) and HEK/TLR4 compared to HEK293/TLR4 + CLI-095 (p=0.0003). In contrast, CLI-095 treatment of HEK293/null did not affect the spread of RSV infection, which was considerably lower than non-treated HEK293/TLR4 controls (Figure 1B).

**Figure 1.**
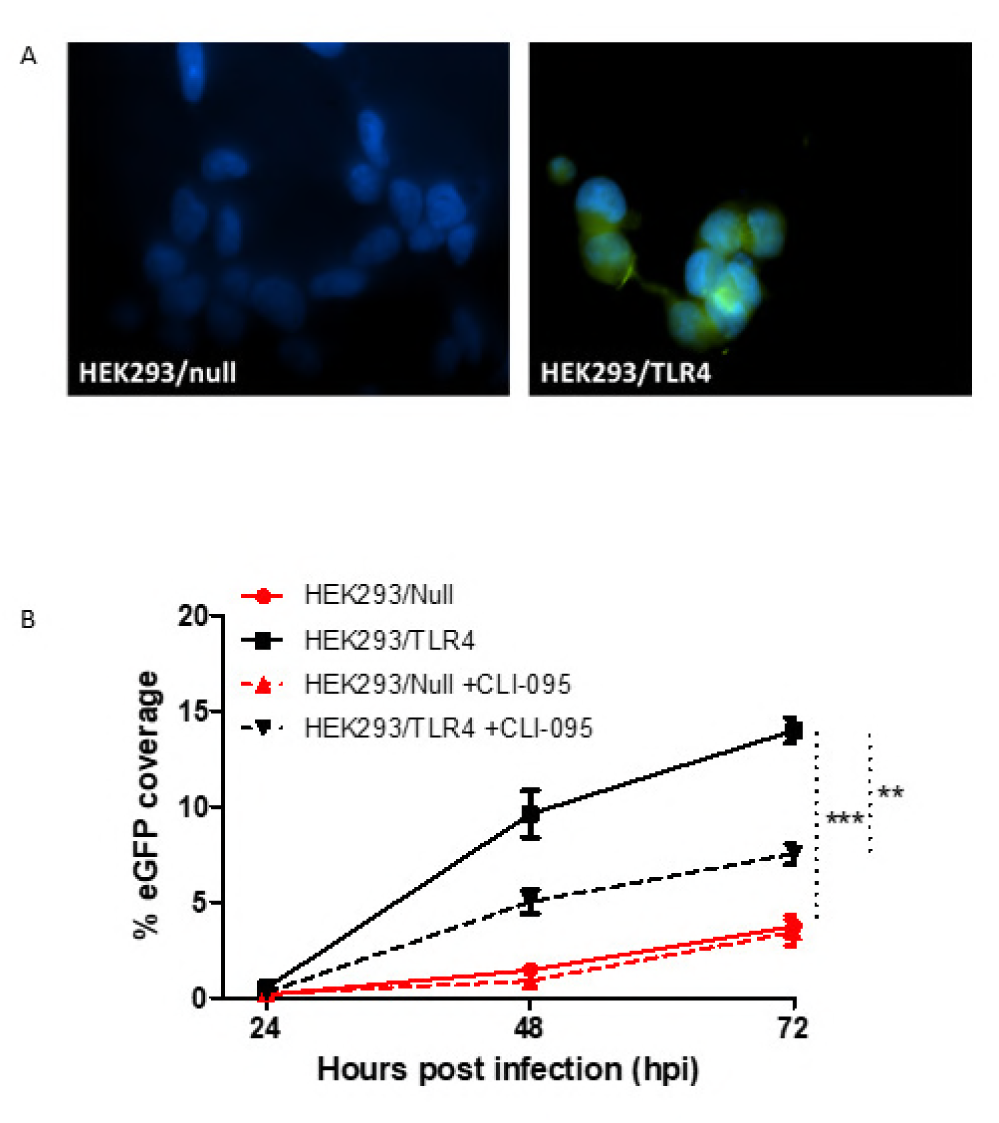
To confirm the presence/absence of TLR4 HEK293 cells stably transfected with TLR4 or null control cells were fixed using 4% PFA (v/v in PBS) then permeabilised using 0.1% Triton X-100 (v/v in PBS) for 1 h. Cells were incubated with anti-TLR4 antibody (Santa Cruz) followed by a green secondary antibody (AlexaFluor) (A). HEK293/null and HEK293/TLR4 cells were treated with 10 μg/mL CLI-095 or mock treated for 6 h then infected (in duplicate) with RSV/eGFP at an MOI=0.1. Five images per well were captured at 24, 48 and 72 hpi using a Nikon TE2000U microscope. The % of the image expressing green fluorescence was assessed using Image J. The average of the 5 fields of view per well was calculated (B). The data were derived from 2 independent experiments carried out in duplicate. Vertical dotted lines indicate statistical significance when areas under the curves were calculated and compared (** p value = <0.01; *** p value = <0.001). Differences were also assessed by t test at each time point.

To confirm whether these data were reproducible in a more physiologically relevant infection model, we exploited our RSV/WD-PBEC model (13,14). WD-PBECs were stained to confirm the presence of TLR4 on the surface of these cells (Figure 2A). WD-PBECs were treated with varying concentrations of CLI-095 prior to infection with a low passage clinical isolate of RSV (RSV BT2a) (MOI=0.1). Consistent with the data from the HEK293/TLR4 cells, a dose-dependent reduction in RSV BT2a growth kinetics was observed (Figure 2B). The two highest concentrations of CLI-095 (10 and 100 μg/mL) resulted in significant reductions in viral titers over the course of the experiment. At the highest CLI-095 concentration used (100 μg/mL) mean RSV titres peaked at 3.09 log_10_ compared to 5.05 log_10_ in untreated controls. At 48, 72 and 96 hpi pre-treatment with 10 or 100 μg/mL CLI-095 resulted in a significant reduction in viral titres compared to the untreated infected controls.

**Figure 2.**
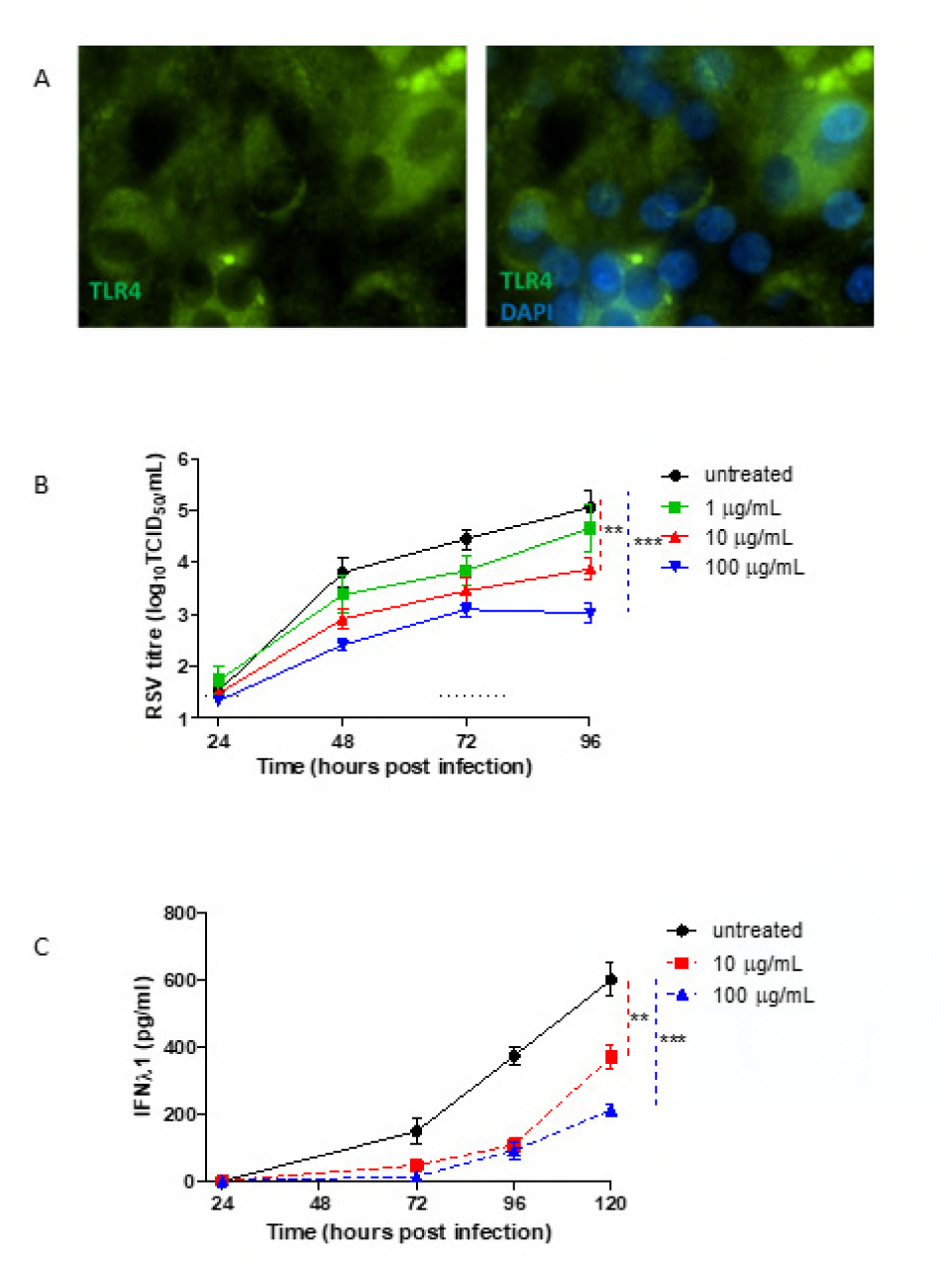
To confirm the presence of TLR4 on WD-PBECs uninfected cultures were fixed using 4% PFA (v/v in PBS) permeabilised then using 0.1% Triton X-100 (v/v in PBS) for 1 h. Cells were incubated with anti-TLR4 antibody (Santa Cruz) followed by a green secondary antibody (AlexaFluor) (A). WD-PBECs (n=3 donors, duplicate Transwells for each condition), pre-treated apically with 1, 10 or 100 μg/mL CLI-095 or untreated, were infected with RSV BT2a (MOI=0.1). Apical washes were harvested every 24 h following infection up to 96 hpi; 200 μL DMEM (with no additives) was incubated on the apical surface for 5 min at RT, removed and snap frozen in liquid nitrogen. Basal medium was also harvested every 24 h. Apical washes were titrated on HEp-2 cells to determine virus growth kinetics (B). IFN-λ1 in basolateral medium from wells, pre-treated with 10 or 100 μg/mL CLI-095 or untreated at 72, 96 and 120 hpi, was quantified by ELISA (eBioscience) (C). Vertical dotted lines indicate statistical significance when areas under the curves were calculated and compared (** p value = <0.01; *** p value = <0.001). Statistical significance between conditions was also assessed at each time point.

Type III IFNs, in particular IFN-λ1, are the main interferons secreted following RSV infection (15). However, the mechanisms by which RSV induces type III IFN expression in epithelial cells is poorly understood. Evidence suggests that TLR4 may be implicated in the induction of type III IFNs following cell stimulation with house dust mite allergens (16). As the RSV F protein was shown to interact with the TLR4/CD14/MD2 complex(8), we hypothesised that type III IFN induction following RSV infection was triggered following activation of the TLR4 pathway. To address this, the basolateral medium from CLI-095-pre-treated RSV BT2a-infected WD-PBECs was harvested from 72 to 120 hpi and the concentration of IFN-λ1 was determined. Treatment with both 10 and 100 μg/mL CLI-095 resulted in substantial reductions in IFN-λ1 secretions (Figure 2C) over the course of the experiment. There was a significant reduction in IFN-λ1 secretion following pre-treatment with 10 or 100 μg/mL CLI-095 at each time point, with the exception of 10 μg/mL at 72 hpi (p=0.0549). However, these CLI-095 concentrations also resulted in significant reductions in virus growth kinetics. If virus replication was essential for IFN-λ1 expression, inhibition of RSV infection/replication, rather than TLR4 signalling per se, might therefore explain the reduction in IFN-λ1 secretion.

To determine if the observed inhibition of RSV growth kinetics or IFN-λ1 production was dependent on specific TLR4 pathway intermediates, small molecule inhibitors directed against specific targets were used. WD-PBECs were treated with inhibitors for PI3K (LY294002), MEK1/2 (U0126), p38 MAPK (SB203580) or NF-κB (JSH-23), or DMSO as a control. Following infection with RSV BT2a, viral titers in apical rinses were quantified every 24 h until 96 hpi (Figure 3A). Inhibition of p38 MAPK resulted in a small but significant decrease in viral growth kinetics compared to the DMSO control. Conversely, MEK1/2 inhibition resulted in a significant increase in viral titers. Basal medium from these experiments was harvested and IFN-λ1 was quantified by ELISA. Significant decreases in IFN-λ1 concentrations were observed following inhibition of either p38MAPK or NF-κB (Figure 3B). In contrast, the significant increases in RSV growth kinetics evident following MEK1/2 inhibition did not result in concomitant increases in IFN-λ1 secretions. Interestingly, the only condition under which the IFN- λ1 concentration was significantly impacted without altering viral titers was NF-κB inhibition, indicating that NF-κB plays an important role in RSV-induced IFN-λ1 induction.

**Figure 3.**
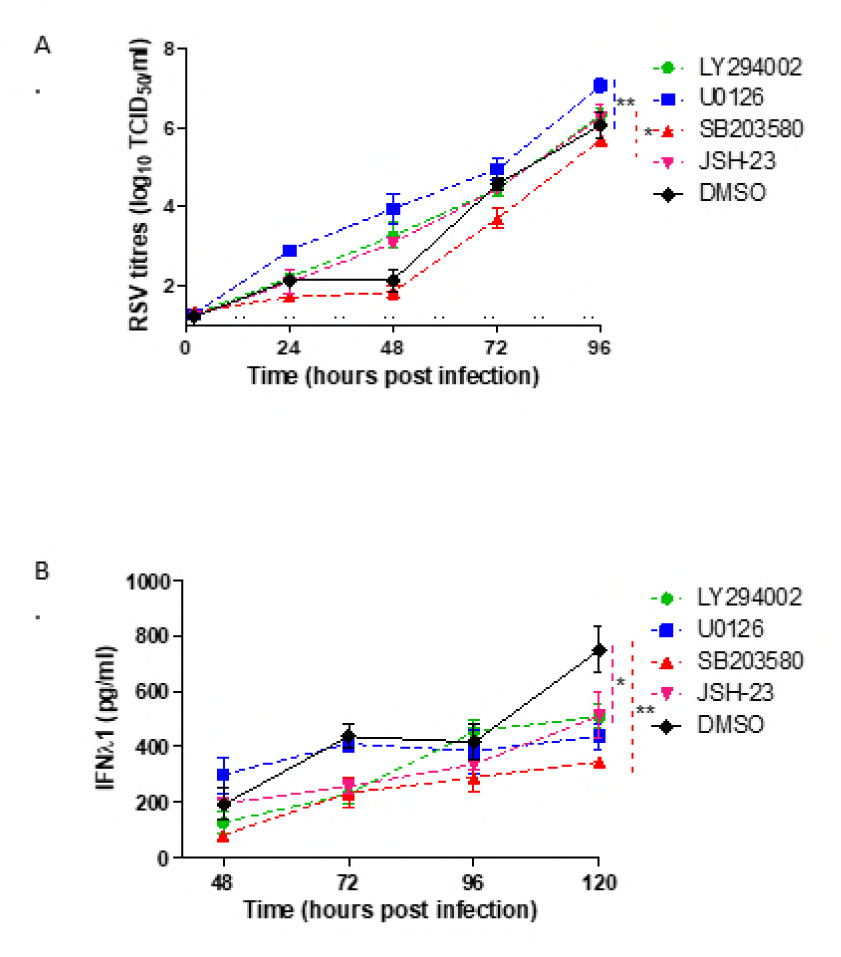
WD-PBECs (n=3 donors) were pre-treated apically with LY294002 (PI3K inhibitor), U0126 (MEK1/2 inhibitor), SB203580 (p38MAPK inhibotor), JSH-23 (NF-κB inhibitor), or DMSO (used as a control) for 1 h at 37° C prior to infection with RSV BT2a (MOI=0.1). Apical washes were harvested every 24 h following infection and titrated on HEp-2 cells to determine virus growth kinetics (A). Basolateral medium was harvested and replaced with fresh medium every 24 h. IFN-λ1 in basolateral medium harvested at 48, 72, 96 and 120 hpi was quantified by ELISA (eBioscience) (B). Vertical dotted lines indicate statistical significance when areas under the curves were calculated and compared * p value = <0.05; ** p value = <0.01).

RSV pathogenesis is mediated in large part by the pro-inflammatory immune responses to infection. Evidence suggests that the production of several chemokines, including IL-6 and CXCL8/IL-8, correlate with the severity of RSV disease in infants(17-20). To determine the consequences of TLR4 inhibition on RSV-induced chemokine secretion levels, WD-PBECs were pre-treated with CLI-095 followed by infection with RSV BT2a. Chemokine concentrations were determined in basal medium harvested at 48, 72 and 96 hpi. CLI-095 treatment prior to RSV infection resulted in significant reductions in MCP-1/CCL2, IL-8/CXCL8, IP-10/CXCL10 and IL- 6 secretions in a dose-dependent manner relative to untreated controls (Figure 4).

**Figure 4.**
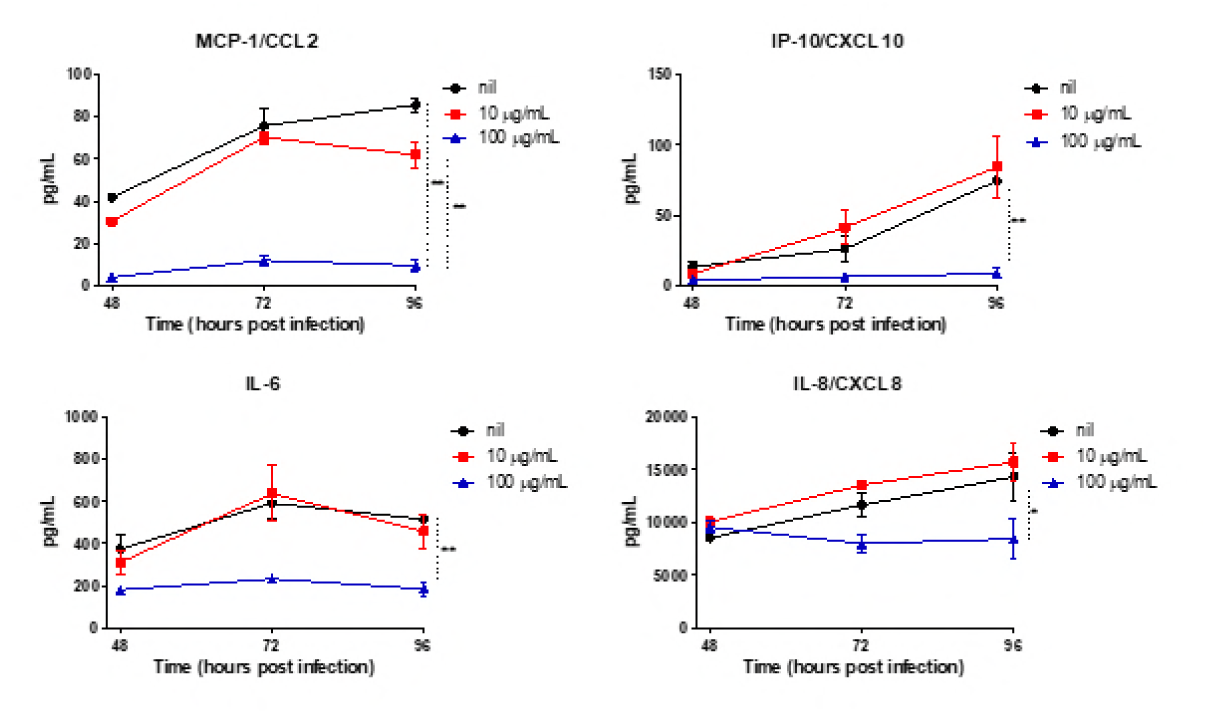
WD-PBECs (n=3 donors), pre-treated apically with 10 or 100 μg/mL CLI-095 or untreated, were infected with RSV BT2a at an MOI=0.1. Basolateral medium was harvested and replaced with fresh medium every 24 h. The concentration of MCP-1/CCL2, IP- 10/CXCL10, IL-6 and IL-8/CXCL8 in basolateral medium was measured by BioPlex (eBioscience) at 48, 72 and 96 hpi. Vertical dotted lines indicate statistical significance when areas under the curves were calculated and compared * p value = <0.05; ** p value =<0.01).

## Discussion

Previous publications reported that TLR4 signalling played no significant role in the entry and/or replication of RSV(8,21). Surprisingly, our data indicated the contrary. We found a significant increase in the level of RSV infection in HEK293/TLR4 cells compared to HEK293/null cells. A number of experimental details may explain this discrepancy with the work of Marr and Turvey (2012)(8). First, while recombinant RSV A2 strains expressing eGFP were used in both studies, the *egfp* gene in our virus was inserted as the 1^st^ transcription unit of the RSV genome, while it was inserted between the RSV *p* and *m* genes in rgRSV. This may have affected the respective virus replication kinetics in the cells. Second, we followed the spread of infection over a longer period of time. In the aforementioned manuscript, infection was followed only until 48 hpi, at which point a trend towards increased RSV spread was evident in HEK293/TLR4 cells compared to HEK/null cells, but it did not reach significance. In contrast, we demonstrated significant differences at both 48 and 72 hpi in these same cell lines. Third, we exploited WD-PBECs and a low passage RSV clinical isolate, RSV BT2a, to confirm the role of TLR4 in RSV infection in a physiologically relevant model. Indeed, we also demonstrated a >2 log_10_ reduction in RSV titers following inhibition of TLR4 intracellular signalling in WD-PBECs with 100 µg/mL CLI-095. In contrast, much of the previous work exploited the use of cell lines that are permissive to RSV infection and can be easily transfected (8). Unlike our RSV/WD-PBEC model, these cell lines do not replicate the morphological or physiological complexities of differentiated airway epithelium, nor the cytopathogenesis of RSV in airway epithelium *in vivo*.

CLI-095 does not interfere with the external domain of TLR4. Therefore, we concluded that the internal portion of the TLR4 complex and/or the downstream signalling is associated with RSV growth kinetics in WD-PBECs. Our data from both immortalised cell lines and WD-PBECs indicated that TLR4 is implicated in RSV infection and/or replication. This is consistent with a recent report demonstrating a protective role of MEG3, a long noncoding RNA, against RSV infection, which most likely acts through the inhibition of the TLR4 signalling pathway(22).

Interestingly, TLR4 antagonists have been associated with a reduction in titers of other viruses both *in vitro* and *in vivo*. Indeed, eritoran, which binds to MD2 and prevents TLR4 activation, decreased viral titers, cytokine production, clinical symptoms and morbidity in a mouse model of influenza virus infection (23).

We demonstrated that CLI-095 possesses substantial prophylactic properties against RSV infection in our WD-PBEC model. As the inhibitor acts by blocking intracellular signalling and does not interfere with the extra-cellular domain of TLR4, it is unlikely that CLI-095 interferes with the RSV F/TLR4 interaction. Indeed, Ii et al demonstrated that CLI-095 does not interfere with the binding of LPS to TLR4(24). As such, diminished RSV growth kinetics are unlikely to be due to the blocking of RSV F protein binding to the TLR4 complex. An association between TLR4 and nucleolin, a putative RSV co-receptor or entry factor, was recently described(25). Further work is needed to establish whether CLI-095 impacted the ability of RSV to interact with nucleolin and thereby, the ability of RSV to infect epithelial cells.

The TLR4 pathway, through adaptor proteins MyD88, TRAM and TRIF, was shown to activate innate immune responses following RSV infection, resulting in the downstream induction of pro-inflammatory cytokines/chemokines.(26) Although significant progress has recently been made in understanding the induction and function of type III IFNs, many questions remain unanswered. It is likely that there is cross-talk between the type I and type III IFN induction pathways, as well as compensatory mechanisms and a degree of redundancy between the two(27). As such, fully elucidating the type III IFN induction pathway has proven complicated. We demonstrated that pre-treatment of RSV-infected WD-PBECs with JSH-23, an NF-κB inhibitor, had a significant impact on the secretion of IFN-λ1 without affecting viral replication. Type III IFN can be induced through both NF-κB-dependent and independent pathways(28). Our data suggest that the induction of IFN-λ1 following RSV infection is, in part, due to NF-κB activation. However, JSH-23 treatment did not completely block IFN-λ1 secretion in response to RSV infection, thereby suggesting that there are other pathways through which RSV triggers IFN-λ1 induction. We also observed a reduction in IFN-λ1 secretion levels following pre-treatment with CLI-095 and SB203580, which were concomitant with significant reductions in viral growth kinetics. It is possible that diminished RSV replication could explain, in part, the decrease in IFN-λ1 secretion. However, the absence of increased IFN-λ1 secretion following MEK1/2 inhibition, despite significantly increased RSV replication following U0126 treatment, suggest that RSV replication kinetics alone do not explain IFN-λ1 secretion levels. Further investigation is required to determine whether the reduction in IFN-λ1 secretion is due to diminished RSV replication or inhibition of TLR4 and/or p38 MAPK signalling, or both.

There is strong evidence to suggest that RSV pathogenesis is immune mediated(29–33). Higher concentrations of IL-6 and IL-8/CXCL8 in stimulated cord blood of infants is predictive of disease severity(19). Furthermore, DeVincenzo *et al*. correlated symptom scores with IL-6 and IL-8/CXCL8 secretion levels and with viral load in a human adult challenge model of RSV disease.(34). Our data demonstrated a reduction in both viral titers and proinflammatory chemokine production following CLI-095 pre-treatment. At the highest CLI-095 concentration used (100 μg/mL) there was a >2 log_10_ reduction in mean RSV titers compared to untreated controls. This reduction in viral replication coincided with a massive reduction in IP-10/CXCL10, MCP-1/CCL2 and IL- 6 secretions and a significant reduction in IL-8/CXCL8. As these chemokines are implicated in RSV pathogenesis, our data are consistent with the concept that restricting RSV replication in infants to a level that poorly stimulates their secretion will alter the disease outcome, thereby providing the rationale for RSV prophylactics or early intervention therapeutics.

In conclusion, intra-cellular signalling mediated by the TLR4 complex was involved in the replication of RSV in WD-PBECs and initiated downstream innate immune responses, including type III IFNs and pro-inflammatory chemokines. Importantly, the TLR4 inhibitor, CLI-095 (or other similar molecules), may have potential as a RSV prophylactic, while our data provides the rationale for exploring its therapeutic potential against RSV in WD-PBECs. p38 MAPK signalling is also implicated in RSV replication. Our data demonstrated that inhibition of TLR4 signalling decreases RSV-induced IFN- λ1 secretion. While the type III IFN induction pathway following RSV infection remains to be fully elucidated, our data demonstrated that both NF-κB and p38 MAPK are implicated. Increasing our understating of the innate immune responses to RSV will aid in the development of RSV prophylactics, therapeutics and vaccines.

## Materials & Methods

### Cell lines and viruses

The origin and characterization of the clinical isolate RSV BT2a were previously described (35). Recombinant RSV expressing eGFP (rRSV A2/eGFP) was a kind gift from Prof. Ralph Tripp (University of Georgia) and Prof. Michael Teng (University of South Florida). Its generation was previously described (36-38). The origin and characterization of the recent clinical isolate RSV BT2a were previously described(39). RSV titers in biological samples were determined as previously described(40). HEK293/TLR4 and HEK293/null cells were obtained from Invivogen and grown in DMEM (4.5 g/L glucose) supplemented with 10% HI-FBS. Cells were kept under selective pressure using blasticidin (Sigma Aldrich). HEK cells were infected for 2 h at 37 °C in DMEM (4.5 g/L glucose, 0% FBS) then maintained in serum-free DMEM (4.5 g/L glucose).

### WD-PBEC cultures

Passage 1 primary paediatric bronchial epithelial cells (PBECs) were obtained commercially (Lonza). WD-PBEC cultures were generated as described previously(14). Complete differentiation took a minimum of 21 days. Cultures were only used when hallmarks of excellent differentiation were evident, including, no holes in the cultures, extensive apical coverage with beating cilia, and obvious mucus production. WD-PBECs were infected apically for 2 h at 37 ° C. For apical rinses of WD-PBECs during experimentation low glucose DMEM was added apically (200 µL) and left for 5 min at room temperature (RT). This was aspirated without damaging the cultures, added to cryovials and snap frozen in liquid nitrogen. At specified intervals post-treatment and/or infection basolateral medium was also harvested and snap frozen in liquid nitrogen, and replaced with fresh medium.

### Immunofluorescence

HEK293/null, HEK293/TLR4 cells and WD-PBECs were fixed with 4% PFA (v/v in PBS) for 40 mins then permeabilised with 0.1% Triton X-100 (v/v in PBS) for 1 h. Cells were blocked with 0.4% BSA (v/v in PBS) for 30 mins then incubated with anti-TLR4 antibody (Santa Cruz) overnight at 4 °C. Following washing, cells were incubated with goat anti-rabbit AlexaFluor 488 antibody for 1 h at 37 °C. Cultures were mounted with DAPI mounting medium (Vectashield, Vector Labs) and imaged using a Nikon Eclipse 90i.

### Signalling inhibitors

All of the inhibitors were reconstituted in DMSO (Sigma Aldrich) as per the manufacturers’ instructions. Concentrations of all inhibitors used for these experiments were at least double the IC_50_(inhibitory concentration 50%) indicated by the manufacturer. Stocks were diluted in ALI medium to achieve the working concentration. WD-PBECs were pre-treated apically at 37 °C for the time indicated in figure legends.

The inhibitors used were:

(i) CLI-095 (Source Bioscience), a TLR4 inhibitor that blocks the intracellular signalling, but not the extracellular domain. Concentrations used are stated in individual figure legends.
(ii) LY294002 (50 µM) (LC labs), a PI3K reversible inhibitor that inhibits Akt/PKB signalling. It also inhibits cell proliferation and induces apoptosis.
(iii) U0126 (20 µM) (LC labs), a highly selective inhibitor of MEK1/2. Acts by disrupting the transcriptional activity of AP-1 and blocks the downstream induction of cytokines and MMPs.
(iv) SB203580 (1 µM) (Sigma Aldrich), a p38 MAPK inhibitor that also blocks PBK phosphorylation and total SAPK/JNK activity. At high concentrations it has been shown to activate the ERK pathway and lead to an increase in NF-κB activity.
(v) JSH-23 (20 µM) (Sigma Aldrich), is an NF-κB inhibitor. It has also been shown, under certain conditions, to inhibit apoptotic chromatin condensation and NO production.

### IFN-λ1 quantification

IFN-λ1 ELISA kits were purchased from eBioscience (Ready to use Platinum Sandwich ELISA kit). IFN-λ1 was quantified from basolateral medium harvested from WD-PBECs, following the manufacturer’s instructions. Frozen aliquots of basolateral medium were rapidly defrosted in a water bath at 37°C and kept on ice during the ELISA procedure to minimise degradation.

### Chemokine quantification

ProcartaPlex kits were purchased from eBioscience to measure a panel of chemokines present in basolateral medium. The manufacturer’s protocol was followed throughout. Analytes measured included IP-10/CXCL10, IL- 8/CXCL8, IL-6, and MCP-1/CCL2.

### Image analysis

Image analysis was carried out using ImageJ software (http://rsbweb.nih.gov/ij/). A minimum of 5 fields were captured per condition/well by UV microscopy (Nikon TE-2000U and Hammamatsu Orca-ER camera).

### Statistical analysis

GraphPad Prism® was used to create graphical representations of the data and for statistical analyses. Summary measures over time were compared by calculating the areas under the curves (AUC) and we used t tests to calculate if these AUCs were statistically significantly different. Differences were also assessed by unadjusted t tests at each time point. p<0.05 was considered significant.

* p value = <0.05; ** p value = <0.01; *** p value = <0.001.

